# Interpretable transcriptome-to-phenotype modeling of cell-painting nuclear morphology features from RNA-seq under low-dose radiation exposure

**DOI:** 10.64898/2026.02.23.707284

**Authors:** Sanket Jantre, Kriti Chopra, Guang Zhao, Clark Cucinell, Rebecca Weinberg, Sara Forrester, Thomas Brettin, Nathan M. Urban, Xiaoning Qian, Byung-Jun Yoon

## Abstract

With rapid advancements in high-throughput multi-modal profiling techniques across molecular, cellular to tissue scales, translating such multi-modal data into knowledge discovery for foundational understanding of cellular mechanisms is central in modern biomedical sciences. In this study, we focus on understanding how low-dose radiation exposure perturbs cellular morphology by linking high-dimensional transcriptomic responses to quantitative cell-phenotype readouts over time. We present a time-resolved inverse modeling framework that associates gene expression changes via RNA-sequencing with nuclear morphology features obtained from cell-painting imaging. Morphology responses were defined as treated-control differences for multiple nuclear features including size, shape, intensity, and textures, indexed by radiation dose and week. To capture time-dependent associations while maintaining interpretability, both RNA-sequencing and cell-painting data were stratified into four temporal phases (weeks 1–2, 3–4, 5–6, 7–9) and phase-dependent effects are encoded via gene-phase interaction predictors. To reduce confounding by dose trends and to evaluate generalization across time, we used a two-stage leave-one-week-out procedure: (i) a dose-only baseline model produced out-of-week residuals for each morphology feature, and (ii) elastic-net regression on phase-aware predictors modeled residual variation not explained by dose. Hyperparameters were selected via an exhaustive grid search scored by the correlation between observed residuals and out-of-week residual predictions, with additional sparsity diagnostics based on nonzero coefficient counts per fold. Stable predictors were identified by selection frequency and sign consistency across folds, then pruned further for multicollinearity and parsimony. Final reduced models were fit using ordinary least squares with heteroskedasticity-consistent standard errors to report effect estimates robust to non-constant variance. This workflow yields a transparent, time-stratified set of transcriptomic predictors associated with longitudinal nuclear morphology changes and provides a reproducible foundation for downstream biological interpretation and validation.

## 1 Introduction

Low-dose ionizing radiation is widely encountered in occupational, environmental, and clinical contexts, yet the molecular-to-phenotypic pathways that connect radiation exposure to measurable cellular morphology perturbations remain incompletely characterized (United Nations Scientific Committee on the Effects of Atomic Radiation (UNSCEAR), 2021). In-vitro longitudinal exposure studies enable systematic measurement of how cellular morphology evolves over time following irradiation, while RNA sequencing (RNA-seq) provides a parallel genome-wide view of transcriptional programs that may underlie these phenotypic changes (Shimabukuro et al., 2022). However, relating high-dimensional gene expression profiles to downstream morphology features is statistically challenging due to limited sample sizes, strong correlation structure among genes, and the possibility that transcriptomic drivers act in a time-dependent manner rather than uniformly across the entire study period.

High-content imaging assays, including Cell Painting (CP) and related nuclear morphology feature sets, offer quantitative descriptors of nuclear size, shape, intensity, and texture (Bray et al., 2016; Cimini et al., 2023). These nuclear features are often sensitive to stress responses, cell-cycle changes, chromatin organization, and remodeling processes that may be induced by radiation exposure (Zheng et al., 2023). Establishing interpretable models that connect RNA-seq to nuclear morphology could therefore serve two complementary purposes: enabling phenotype prediction from molecular readouts and generating mechanistic hypotheses about which transcriptional programs align with early versus late morphology changes.

However, directly regressing morphology on thousands of gene expression measurements is statistically troublesome due to being prone to overfitting and confounding. First, both gene expression and morphology often exhibit strong week-to-week time structure that can confound transcriptome–phenotype associations, and morphology outcomes may be dominated by dose-related trends (Wang et al., 2025). Second, gene expression predictors are highly correlated, making ordinary least squares (OLS) regression-based coefficient estimates unstable and difficult to interpret without regularization (Hastie et al., 2009). Third, it is plausible that different transcriptomic drivers act at different stages of the response trajectory in a time-dependent manner rather than uniformly across the entire study, motivating models that allow time-local (or phase-local) associations rather than a single global effect shared across all weeks.

Accordingly, there is a need for modeling strategies that (i) isolate the variance in morphology that is not trivially explained by simple dose structure, (ii) perform principled variable selection in a high-dimensional setting, (iii) provide interpretable, phase-aware predictors, and (iv) evaluate generalization in a manner aligned with the longitudinal design (e.g., leaving out entire weeks rather than random samples).

In this work, we develop an interpretable transcriptome-to-phenotype modeling pipeline that links RNA-seq responses to multiple nuclear morphology feature changes measured across weeks under low-dose radiation exposure. The central idea is to separate baseline dose-related structure from residual variation that may be explained by transcriptomic signals, while enforcing interpretability through phase-dependent gene effects. We construct morphology outcomes as differences relative to controls at each week and compute gene expression responses as within-week log_2_-fold-changes relative to baseline. Next, we define a set of candidate genes by aggregating highly responsive genes per week across weeks and assigning them to non-overlapping temporal phases. We then create phase-aware gene predictors by interacting gene responses with phase indicators, allowing a gene to contribute differently depending on the stage of the longitudinal response.

In order to evaluate generalization in a manner aligned with the longitudinal design, we use leave-one-week-out (LOWO) cross-validation. We first fit a dose-only baseline model within each held-out week and compute residuals, then fit sparse regularized models based on Elastic Net regression (Zou and Hastie, 2005) on these residuals within the same cross-validation splits, sweeping hyperparameters to identify settings that best predict residual variation. For morphology features where residuals are meaningfully predictable from transcriptomic predictors, we rank and shortlist phase-local gene predictors by selection stability and sign consistency across held-out weeks and further reduce them based on multicollinearity and parsimony checks. At last, we fit a final reduced regression model using OLS with heteroskedasticity-consistent (HC3) standard errors to report interpretable effect sizes along with their confidence intervals (Long and Ervin, 2000).

This design yields two complementary outputs: (i) predictive performance estimates under a week-held-out evaluation that matches the longitudinal structure, and (ii) an interpretable set of phase-dependent gene associations that can be examined as candidates for biological follow-up studies.

The primary contributions of this study are:

- A modeling framework for relating RNA-seq responses to nuclear morphology features in a longitudinal low-dose radiation exposure setting, with validation and evaluation based on leaving out entire weeks.
- A phase-aware representation of transcriptomic predictors that enables time-dependent interpretability while controlling for dose-driven baseline structure.
- A robustness-focused gene selection strategy combining regularized residual modeling, hyperparameter sweeps, and stability-based ranking, followed by interpretable regression with compact set of phase-dependent gene effects.

We emphasize that the resulting gene–phase associations are intended to be predictive and hypothesis-generating rather than definitive mechanistic biological claims. Establishing causal biological relevance and mapping these predictors to known pathways will require additional domain-informed analysis and validation, which we consider a key direction for future work.

## 2 Methodology

### 2.1 Study design and data modalities

We analyze a longitudinal in-vitro low-dose radiation exposure study with measurements collected across multiple discrete time-points (weeks) and dose conditions. For each experimental unit (e.g., replicate well or sample) we consider: (i) transcriptomic measurements from RNA sequencing (RNA-seq), and (ii) high-content imaging-derived nuclear morphology features computed from cell painting images which undergo automated nucleus segmentation using a Cell-SAM-based pipeline (Marks et al., 2025). The longitudinal structure induces strong week-to-week shifts in both gene expression and morphology; therefore, our modeling and validation are designed around holding out entire weeks rather than randomly splitting samples.

#### 2.1.1 Morphology outcomes and normalization

We extract both structural and texture nuclear features from Cell Painting images using the DNA channel stained with Hoechst 33342 (Roach et al., 2026). Nuclei are first segmented from the Hoechst channel using Cell-SAM to obtain instance-level masks, from which feature measurements are computed for individual nucleus. For each week-dose condition, feature values are averaged across thousands of segmented nuclei, yielding a condition-level summary. Let *y*_*w,d,f*_ denote the condition-level summary of nuclear morphology feature *f* measured at week *w* and dose condition *d*. Next, to reduce nuisance variation and focus on exposure-related changes, we express morphology outcomes as within-week differences relative to matched controls. Specifically, for each week *w* and non-control dose *d*, we compute

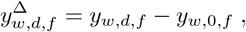

where *y*_*w*,0,*f*_ denotes the feature value under the control condition measured in the same week. This “difference-from-control” transformation emphasizes perturbation magnitude and improves comparability across weeks by absorbing week-specific baseline effects. Stacking all non-control (*w, d*) conditions yields the outcome matrix *Y* ∈ ℝ^*n*×*F*^, where *n* is the number of week–dose samples (excluding controls) and *F* is the number of morphology features.

#### 2.1.2 RNA-seq processing and gene response definition

Raw RNA-seq reads were preprocessed by the sequencing lab at University of Illinois Urbana-Champaign, which removed all low-quality sequences from the data. Trimmed reads were aligned to the human reference genome GRCh38.p14 (NCBI RefSeq assembly GCF_000001405.40; RefSeq annotation RS_2023_10, 2023-10-02) using STAR v2.7.11b with `--runThreadN 16`, reading compressed inputs via `--readFilesCommand zcat` (paired-end), and emitting coordinate-sorted BAM files (`--outSAMtype BAM SortedByCoordinate`) with `--outSAMstrandField intronMotif`. BAM files were indexed using samtools v1.20. Gene-level read counts were quantified from aligned BAMs using featureCounts v2.0.6 with exon-level counting and gene-level summarization (`-t exon -g gene id`), using -p for paired-end fragment counting and 16 threads (`-T 16`); transcripts-per-million (TPM) values were additionally computed per sample using TPMCalcu-lator v0.0.3 (`-e -a`; paired-end: -p). Differential expression analysis was performed using pyDESeq2 v0.4.9 with a one-factor design (`design factors = ‘Condition’`) and Cook’s-distance refitting enabled (`refit_cooks=True`). For each user-specified week-matched dose-versus-control contrast, Wald tests were performed via `contrast = [‘Condition’, cond1, cond2]` and *p*-values were adjusted for multiple testing using the Benjamini–Hochberg false discovery rate (FDR). Genes with adjusted *p*-value < 0.05 and |log_2_ FoldChange| > 1 were considered differentially expressed and used downstream for week-specific gene screening and construction of gene predictors.

RNA-seq measurements are processed into gene-level responses intended to capture exposure-associated transcriptional changes within each week. Let *x*_*w,d,g*_ denote the normalized expression level (or abundance) of gene *g* measured at week *w* and dose condition *d*, after standard preprocessing (e.g., alignment/quantification, filtering, normalization). To align predictors with the difference-from-control morphology outcomes and reduce week-to-week nuisance shifts, we define within-week gene responses relative to matched controls. For each week *w*, non-control dose *d*, and gene *g*, we compute a response such as a log_2_-fold-change

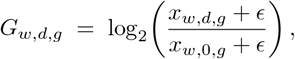

where *x*_*w*,0,*g*_ is the same-week control measurement and *ϵ >* 0 is a small offset to stabilize ratios for low-expression genes. Alternative within-week response definitions (e.g., centered differences on a transformed scale) are compatible with the pipeline and can be substituted without changing downstream modeling. Stacking all non-control (*w, d*) pairs yields the gene response matrix *G* ∈ ℝ^*n*×*P*^, where *n* is the number of week–dose samples (excluding controls) and *P* is the number of genes retained for analysis.

#### 2.1.3 Candidate gene filtering and phase construction

Directly modeling genome-wide expression across thousands of genes is statistically unstable in small-*n* settings, particularly under strong gene–gene correlations and week-to-week shifts. We therefore construct a compact, response-enriched candidate gene set via a screening procedure that preserves the longitudinal structure and avoids pooling across weeks in a way that could blur time-local effects.

For each week *w*, we identify the top *K* most responsive genes across all non-control dose conditions using week-stratified differential expression results. Concretely, genes are ordered by statistical significance (by p-values) from the within-week contrasts comparing each dose to the same-week control, and the top *K* genes are retained for that week. To explicitly model time-local associations, we partition the longitudinal timeline into a small number of non-overlapping temporal phases corresponding to contiguous week ranges. In the current analysis, we use four phases motivated by an integrated qualitative summary of the chronic low-dose-rate response trajectory (Figure 2): weeks 1–2 (“alarm”), 3–4 (“remodel”), 5–6 (“integrate”), and 7–9 (“rewire”). The phase names are shorthand labels for early-to-late response intervals and are not intended to assert pathway identity. We define a phase-specific candidate gene set by taking the union of week-level responsive genes within that phase,

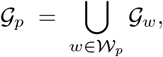

where 𝒢_*w*_ is the set of top-*K* responsive genes in week *w* and 𝒲_*p*_ is the set of weeks assigned to phase *p*. This yields phase-specific candidate pools, enabling the downstream model to assign a gene a nonzero effect in some phases while allowing it to be inactive in others. In the current study we set *K* = 1000 (per week) to retain a broad, response-enriched search space; taking unions within each phase results in candidate set sizes |𝒢_alarm_| = 1368, |𝒢_remodel_| = 1699,|𝒢_integrate_| = 1571, and |𝒢_rewire_| = 1959. Candidate gene screening is performed once using all weeks and treated as a fixed feature-engineering step. Accordingly, the reported LOWO predictive metrics should be interpreted as conditional on this globally screened candidate set, rather than as fully nested performance under fold-specific screening.

**Figure 1.**
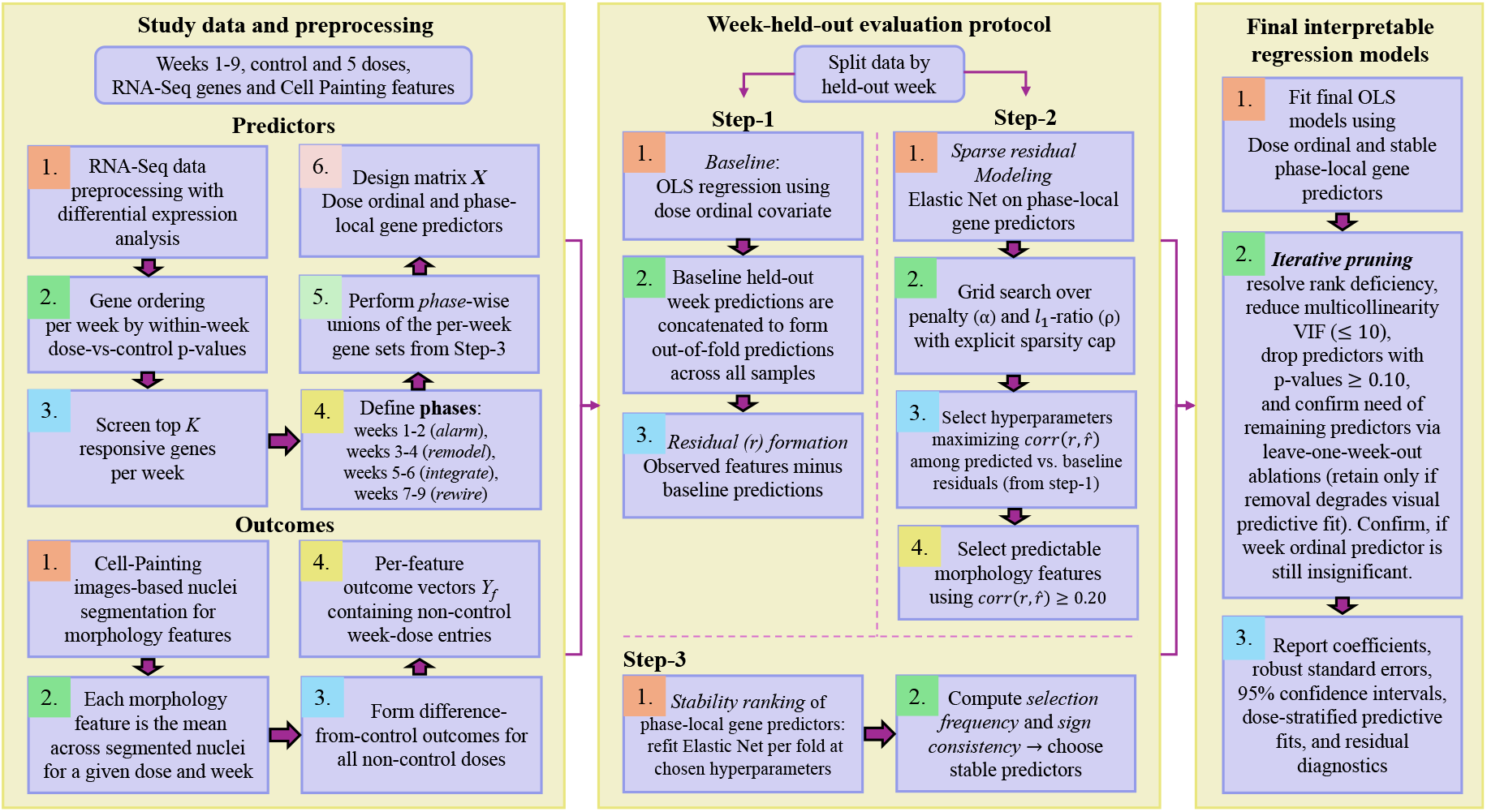
Workflow schematic for the transcriptome-to-phenotype modeling pipeline under leave-one-week-out (LOWO) evaluation. RNA-seq differential-expression results are used to screen week-specific responsive genes, which are then pooled within four temporal phases to construct phase-aware gene predictors in the design matrix. Morphology responses are computed from cell-painting nuclear features as within-week differences from matched controls. Modeling proceeds with a dose-only baseline (Step 1), Elastic Net regression-based sparse residual modeling with LOWO hyperparameter selection (Step 2), stability ranking of phase-aware predictors across folds (Step 3), and a final pruned ordinary-least-squares (OLS) regression model for interpretable effect estimation.

**Figure 2.**
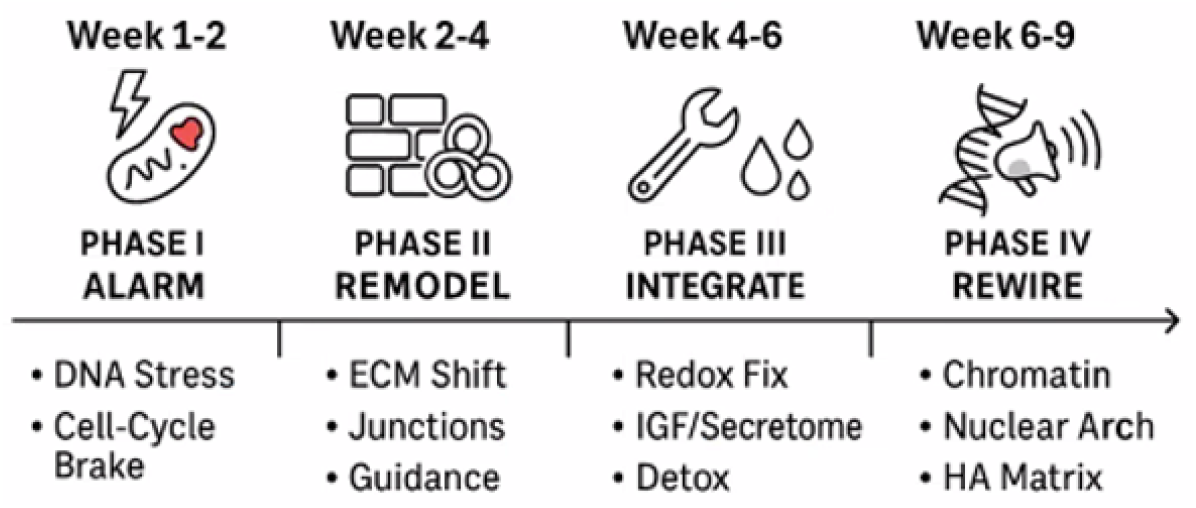
Temporal phase partition used throughout the analysis. Weeks are grouped into four contiguous phases–weeks 1–2 (alarm), 3–4 (remodel), 5–6 (integrate), and 7–9 (rewire)–to define phase-specific candidate gene pools and to construct phase-local gene predictors via interactions between gene responses and phase indicators. Phase names are shorthand labels for early-to-late response intervals and do not imply specific pathway identity.

Finally, the specific four-phase segmentation and labels were refined using an AI-assisted hypothesis-generation prompt given to a large language model, which was used only to propose a coherent temporal narrative for grouping contiguous weeks. All statistical selection, modeling, and validation steps are performed on the measured data as described in the subsequent sections.

#### 2.1.4 Design matrix construction

Each observation corresponds to a week–dose condition (*w, d*) (excluding control), where *w* ∈ {1, …, 9} denotes week and *d* ∈ {*F, G, H, I, J*} denotes a non-control dose level. After aligning the RNA-seq response table and the morphology outcome table to the common set of week–dose conditions, we construct a single predictor matrix *X* with one row per (*w, d*).

##### Baseline exposure covariate

We encode dose as an ordinal covariate, `dose_ord` (*d*) ∈ {1, 2, …, 5}, using a fixed mapping from dose labels to integers (*F* → 1, *G* → 2, ···, *J*→ 5). This parsimonious baseline predictor captures dominant monotone exposure structure while remaining interpretable.

##### Phase-local gene effects

Using the phase partition defined in Section 2.1.3 (Figure 2), let 𝕀{`phase` (*w*) = *p*} denote the indicator for phase *p*. Let *G*_*w,d,g*_ denote the within-week gene response (log_2_-fold-change versus the same-week control) for gene *g* at condition (*w, d*). Given phase-specific candidate sets 𝒢_*p*_, we define phase-local predictors via gene–phase interactions:

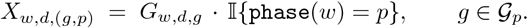

This construction enforces interpretability: each coefficient corresponds to the association between a gene response and a morphology feature within a specific temporal phase. By design, *X*_*w,d*,(*g,p*)_ = 0 outside phase *p*, preventing leakage of phase-local gene effects across the timeline.

##### Handling missingness

After constructing all derived predictors, any remaining missing values (arising from alignment or upstream preprocessing) are set to zero. In practice, because interaction predictors are structurally zero outside their phase and only defined for available gene responses, this step acts primarily as a conservative safeguard against sparse missing entries rather than as substantive imputation.

### 2.2 Week-held-out evaluation protocol

Because week-to-week shifts can dominate both data modalities, we evaluate generalization using leave-one-week-out cross-validation (LOWO). Let 𝒲 be the set of weeks. For each held-out week *w* ∈ 𝒲, the training set contains all samples not in week *w*, and the test set contains all samples from week *w*. Predictions are concatenated across folds to produce out-of-fold predictions for all samples. Unless stated otherwise, all model fitting, scaling, hyperparameter tuning, and feature ranking are performed within this LOWO framework.

#### 2.2.1 Step 1: baseline dose-only model

For each morphology feature *f*, we first fit a baseline model that captures the dominant dose-driven trend without using transcriptomic predictors. Within each LOWO fold, we fit an ordinary least squares (OLS) regression on the training data with intercept and `dose_ord`:

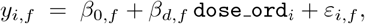

and generate test-week predictions 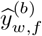. We then define residuals

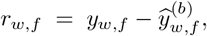

which represent the portion of feature *f* not explained by the dose-only baseline under week-held-out prediction. This residual formation step is crucial: it forces the subsequent transcriptomic model to explain variation beyond what dose alone predicts, rather than rediscovering dose structure through correlated gene responses.

#### 2.2.2 Step 2: sparse residual modeling

To test whether phase-local gene effects can explain residual variation, we fit a sparse regression model mapping phase-local gene predictors to residuals. Within each LOWO fold, we standardize gene predictors using a training-fold-based standard normal transform (z-score) and apply the same scaling to the held-out week. We then fit Elastic Net regression (Zou and Hastie, 2005) without intercept (because we explicitly center the training residuals and add the mean back at prediction time):

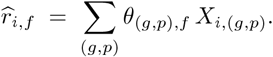

Elastic Net combines *ℓ*_1_ and *ℓ*_2_ penalties, enabling sparse selection while mitigating instability under correlated predictors. For each morphology feature *f*, we sweep a grid of hyperparameters (*α, ρ*), where *α* is the overall penalty strength and *ρ* is the *ℓ*_1_ ratio (with *ρ* = 0 and 1 corresponding to Ridge (Hoerl and Kennard, 1970) and Lasso (Tibshirani, 1996) regression respectively). For each grid setting we compute out-of-fold predictions 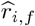 and score performance using the correlation 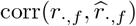. This metric directly quantifies how well gene predictors recover residual structure that the baseline dose-only model leaves unexplained.

##### Grid sweep with optimized computation

A naive grid sweep refits Elastic Net for every (*α, ρ*) within every fold, which can be slow. To accelerate, we use the coordinate descent path solver (e.g., `enet_path`) to compute solutions for a decreasing sequence of *α* values in one call per fold and *ρ*, reusing warm starts. This yields predictions for all *α* at fixed *ρ* with a single path computation per fold.

##### Sparsity control

As downstream stability ranking and interpretability depend on sparse solutions, we track the number of nonzero coefficients (*nnz*) per fold for each (*α, ρ*). This allows us to enforce a sparsity cap (e.g., maximum *nnz* per fold ≤ 50) when selecting the best hyperparameters–(*α, ρ*), choosing the highest-scoring setting among those that satisfy the cap. If no grid setting satisfies the cap, we fall back to the overall best-scoring setting. This mechanism prevents selecting hyperparameters that achieve higher residual correlation only by fitting dense, less interpretable models.

Implementation details of the Elastic Net regression grid search (values of *α* and *ρ*), centering/standardization, and solver settings are provided in Appendix A.

#### 2.2.3 Step 3: stability ranking of phase-local gene effects

For each morphology feature that shows meaningful residual predictability (e.g., 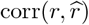) above a threshold of ≥ 0.20), we quantify the stability of selected phase-local gene effects across LOWO folds. Fixing the chosen (*α, ρ*) for that feature, we refit Elastic Net regression within each training fold and record which coefficients are nonzero. For each predictor *t* = (*g, p*) we compute the selection frequency

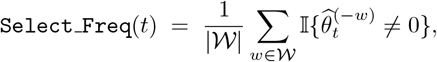

and the sign-consistency score

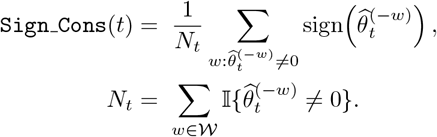

*N*_*t*_ indicates the number of folds in which the predictor is selected. `Sign_Cons` (*t*) ∈ − [1, 1] indicates whether the direction of association is consistent across folds. Predictors are ranked primarily by `Select_Freq` (*t*) (descending) and secondarily by a deterministic tie-breaker (e.g., lexicographic order of the predictor name) to ensure reproducibility. In our analysis, the top-ranked predictors typically exhibit highly consistent directions (often near *±* 1), so ranking is driven mainly by selection frequency. Overall, this yields a compact set of phase-local gene effects that are repeatedly supported across LOWO fits.

### 2.3 Final interpretable regression and inference

To report interpretable phase-local gene effects, we fit a final parsimonious linear model for each selected morphology feature using ordinary least squares (OLS) with heteroskedasticity-consistent (HC3) standard errors (Long and Ervin, 2000). The final model includes an ordinal dose covariate and a subset of stable phase-local gene predictors:

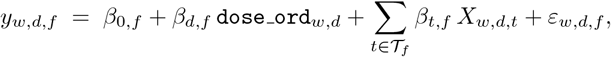

where 𝒯_*f*_ is determined from the stability ranking (e.g., retaining predictors with `Select_Freq` ≥ 7/9). HC3 standard errors are used to reduce sensitivity to heteroskedasticity and leverage effects that can be pronounced in small-sample regression settings.

#### 2.3.1 Model pruning and multicollinearity checks

Because phase-local predictors can be highly correlated, we apply an explicit pruning procedure before reporting final OLS coefficients. Multicollinearity is assessed using variance inflation factors (VIF). When the design matrix is rank-deficient, VIF values can become infinite due to exact (or near-exact) linear dependencies among predictors. In such cases, we first restore full rank by removing one predictor that contributes most strongly to a dependency direction identified via singular value decomposition (SVD) of the centered design matrix. This step is repeated until the matrix is full-rank and all VIF values are finite.

After resolving rank deficiency, we iteratively remove predictors with the largest VIF until all remaining predictors satisfy a conservative multicollinearity threshold (VIF *<* 10). We then perform additional pruning based on statistical evidence and parsimony: predictors with weak evidence of association (e.g., *p >* 0.10 under HC3 standard errors) are removed one at a time until all retained predictors meet the chosen statistical significance threshold.

Finally, we validate the necessity of each remaining predictor—including the ordinal dose covariate— using week-held-out predictive performance. Specifically, we compare leave-one-week-out (LOWO) *R*^2^ for models fit with and without a candidate predictor and assess whether its removal causes a material decrease in out-of-week generalization. We additionally monitor in-sample fit summaries (e.g., *R*^2^ and adjusted *R*^2^) to ensure that pruning does not substantially degrade the qualitative model behavior. In practice, only a small number of pruning iterations (typically ≤ 3–4) are performed to avoid over-pruning at this stage.

As a final parsimony check, we reassess an auxiliary week-ordinal covariate. If it is not statistically supported after including the dose-ordinal covariate and selected phase-local gene effects (e.g., *p >* 0.10), we exclude it; this is consistent with the interpretation that residual time-dependent variability (beyond the baseline dose trend) is captured by the phase-local transcriptomic predictors. We also record whether the dose-ordinal predictor remains supported after adding transcriptomic predictors, treating this as a diagnostic of how much dose-associated variation is absorbed by the selected gene effects rather than as a criterion for inclusion.

##### Reporting and reproducibility

For each morphology feature, we report: (i) the baseline dose-only LOWO performance, (ii) residual predictability from phase-local gene effects under Elastic Net with selected hyperparameters, (iii) the stability-ranked set of phase-local predictors with selection frequency and sign consistency, and (iv) final OLS(HC3) coefficient estimates with confidence intervals. All computations are implemented in Python using standard scientific libraries (e.g., `numpy, pandas, scikit-learn, statsmodels`), and evaluation is deterministic given fixed inputs. Where iterative solvers are used, convergence criteria (maximum iterations and tolerance) are set conservatively; warnings are monitored and, when necessary, the regularization grid is expanded toward stronger penalties to ensure stable solutions.

## 3 Results

### 3.1 Data preprocessing

#### 3.1.1 Difference-from-control normalization of morphology features

We first assessed nuclear morphology features before and after applying the within-week difference-from-control transformation (Section 2). In the raw feature space, many endpoints display pronounced week-to-week shifts in baseline levels that are not purely attributable to dose, and in several cases are comparable in magnitude to the dose-associated changes themselves (Figure 3). Such baseline drift is consistent with nuisance variation in longitudinal in-vitro assays (e.g., culture and passage effects, imaging and segmentation variability, or other week-specific factors) and motivates evaluation protocols that explicitly hold out entire weeks.

**Figure 3.**
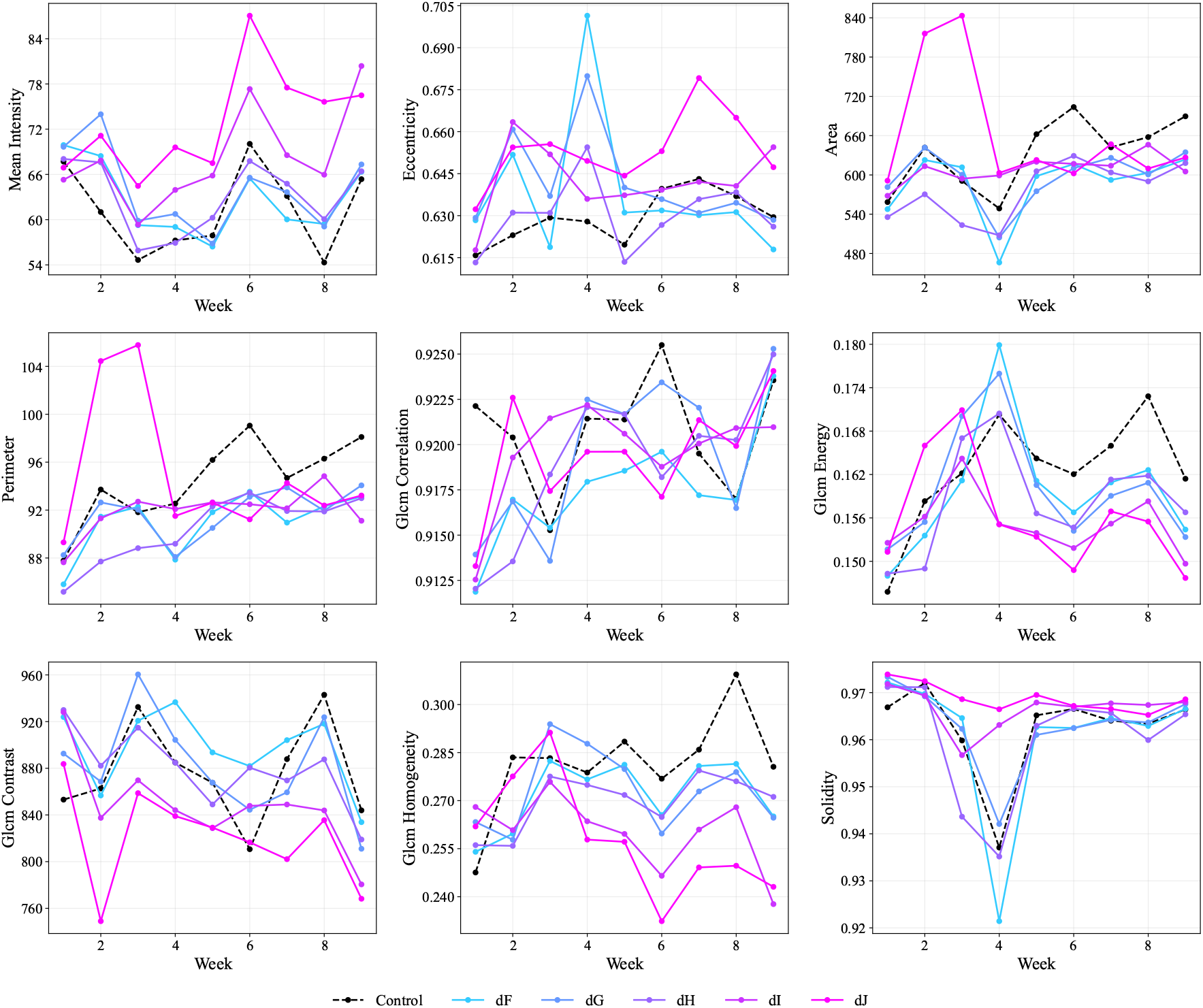
Overview of raw nuclear morphology feature trajectories across weeks and dose levels. Each panel shows the mean value of a single feature (area, perimeter, eccentricity, solidity, mean intensity, and selected GLCM texture features) for each week–dose condition. Colored lines denote non-control dose levels (dF–dJ), while the control condition is shown as a dashed black line with markers. This figure summarizes the dominant week-to-week structure and dose-dependent trends prior to within-week normalization.

Applying the difference-from-control normalization reduces these week-specific baselines by anchoring each week to its matched control condition. The resulting Δ-features are identically zero for controls by construction and more directly reflect radiation exposure-associated deviations within each week (Figure 4). This transformation improves comparability across weeks and emphasizes deviations that are plausibly attributable to exposure rather than global offsets. Given the strong longitudinal structure apparent in the raw measurements, all subsequent modeling results are reported under leave-one-week-out (LOWO) cross-validation, which tests whether models generalize to unseen weeks rather than leveraging shared within-week structure.

**Figure 4.**
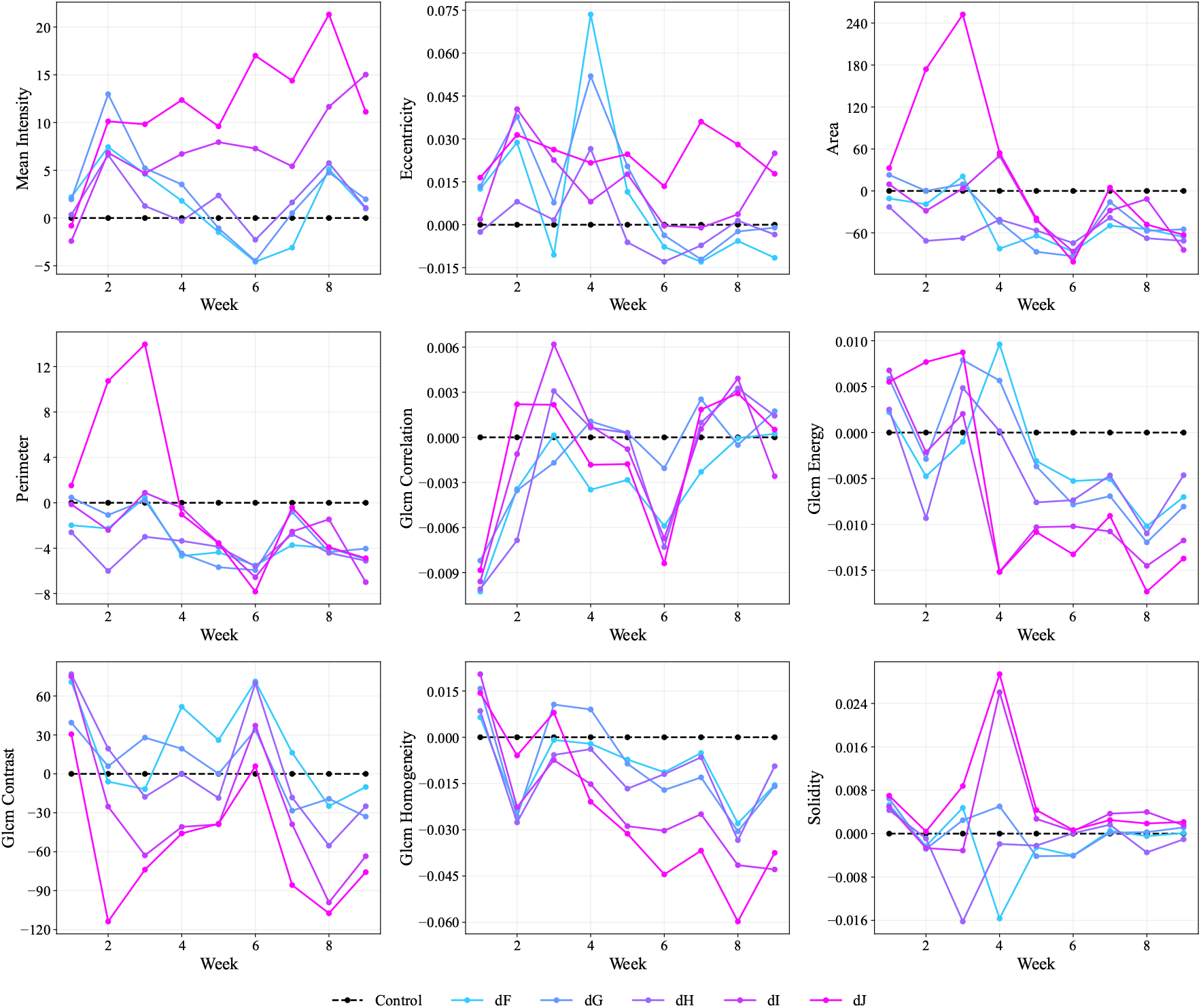
Overview of within-week difference-from-control nuclear morphology trajectories. Each panel shows Δfeature(*w, d*) = feature(*w, d*) − feature(*w*, 0) across weeks for each non-control dose level (dF–dJ). The control condition is included and is identically zero (dashed black line with markers), providing a visual reference for the normalization. This representation removes week-specific baselines and highlights exposure-associated deviations relative to matched same-week controls.

### 3.2 Week-held-out evaluations

#### 3.2.1 Dose-only baseline and residual modeling

For each morphology feature *f*, we first fit a dose-only baseline model under leave-one-week-out (LOWO) cross-validation and form week-held-out residuals *r*_·,*f*_ (Section 2). This baseline provides a stringent reference point: any downstream transcriptome-based improvement must explain variation that generalizes to an entirely unseen week, rather than rediscovering dose structure indirectly through correlated gene responses. We summarize baseline LOWO performance together with residual-model performance in Table 1.

**Table 1:** Baseline vs. residual predictability across nuclear morphology features. For each feature, we report leave-one-week-out (LOWO) performance of the dose-only baseline model (LOWO *R*^2^ and correlation), residual predictability from phase-local transcriptomic predictors measured as 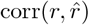, the selected Elastic Net hyperparameters (*α, ρ*), and sparsity as the median and maximum number of nonzero coefficients (nnz) across the 9 folds.

| Morphology feature | Baseline LOWO $R^2$ | Baseline LOWO corr | corr($r, \hat{r}$) | $\alpha$ | $\rho$ | Median nnz/fold | Max nnz/fold |
| --- | --- | --- | --- | --- | --- | --- | --- |
| mean intensity | 0.233 | 0.494 | 0.451 | 0.10 | 0.95 | 44 | 49 |
| eccentricity | -0.140 | -0.137 | 0.371 | 0.001 | 0.80 | 30 | 31 |
| area | -0.014 | 0.162 | 0.346 | 1.00 | 1.00 | 33 | 38 |
| perimeter | -0.068 | 0.046 | 0.312 | 0.03 | 1.00 | 36 | 42 |
| glcm correlation | -0.204 | -0.498 | 0.268 | 0.001 | 0.80 | 15 | 17 |
| glcm energy | -0.171 | -0.198 | 0.151 | 0.001 | 0.80 | — | — |
| solidity | -0.017 | 0.172 | 0.120 | 0.003 | 1.00 | — | — |
| glcm contrast | 0.130 | 0.392 | -0.026 | 10.0 | 1.00 | — | — |
| glcm homogeneity | -0.101 | 0.028 | -0.102 | 0.001 | 1.00 | — | — |

Baseline performance varies substantially across features. Several features exhibit negative LOWO *R*^2^, which indicates that, when evaluated on held-out weeks, the dose-only predictor performs worse than a trivial predictor that always outputs the training-fold mean. Negative *R*^2^ in this setting is not unusual: it reflects distribution shift across weeks and the fact that a single monotone dose predictor cannot capture week-specific offsets or nonstationary exposure responses. The baseline LOWO correlation provides a complementary summary of directional agreement between predicted and observed values; negative correlations indicate systematic sign mismatch on held-out weeks, whereas small positive correlations can coexist with negative *R*^2^ when predictions capture the direction but not the scale (or are poorly calibrated across weeks).

These observations motivate our two-stage residual modeling strategy. By defining *r*_·,*f*_ using week-held-out baseline predictions, we ensure that the subsequent Elastic Net model is tasked specifically with explaining the structured variance that remains after accounting for the dose-only trend under the same LOWO splits. In this way, positive 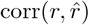 reflects transcriptome-based recovery of residual structure that generalizes across weeks, while avoiding optimistic estimates that could arise if residuals were computed in-sample.

#### 3.2.2 Sparse phase-local transcriptomic models predict residual variation for five features

We next tested whether phase-local transcriptomic predictors explain morphology variation beyond the monotone dose trend captured by the baseline model. Specifically, for each feature *f* we fit Elastic Net regression models under LOWO to predict the baseline residuals from phase-local gene predictors, selecting (*α, ρ*) to maximize 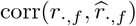 while enforcing sparsity through an upper bound on the number of nonzero coefficients per fold (Section 2). This setup ensures that improved performance reflects recovery of structured residual signal rather than rediscovery of dose effects through correlated transcriptomic responses.

Residual predictability was concentrated in five out of nine nuclear morphology features. Using 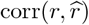) ≥ 0.20 as an operational threshold, five features showed meaningful week-held-out residual predictability: `mean intensity, eccentricity, area, perimeter`, and `glcm correlation` (Table 1). For these features, the selected hyperparameters produced sparse solutions with median nnz per fold ranging from ≈ 15 to ≈ 44 (and fold-wise maximum remaining modest), supporting the goal of interpretability-oriented downstream analysis. In contrast, the remaining morphology features exhibited weak or negative residual correlations, suggesting that after removing the dose-only baseline, their remaining variation is not reliably predictable from the current phase-local transcriptomic representation under week-held-out evaluation.

#### 3.2.3 Stability ranking yields compact sets of phase-local gene predictors

For each nuclear morphology feature exceeding the residual predictability threshold, we refit Elastic Net regression within each LOWO training fold at the selected (*α, ρ*) and quantify stability of phase-local gene predictors using selection frequency and sign consistency (Section 2). Using a strict stability threshold of `Select_Freq` ≥ 7/9 (selected in at least 7/9 folds) yielded compact sets of predictors, ranging from 10 to 32 phase-local gene predictors per morphology feature (Table 2). Predictors are ordered primarily by selection frequency and, for ties, by lexicographic order. Across features, the retained predictors exhibited strongly consistent directions (sign consistency of *±* 1 for all retained predictors), indicating that when a predictor was selected, the direction of its estimated association was stable across folds.

**Table 2:**
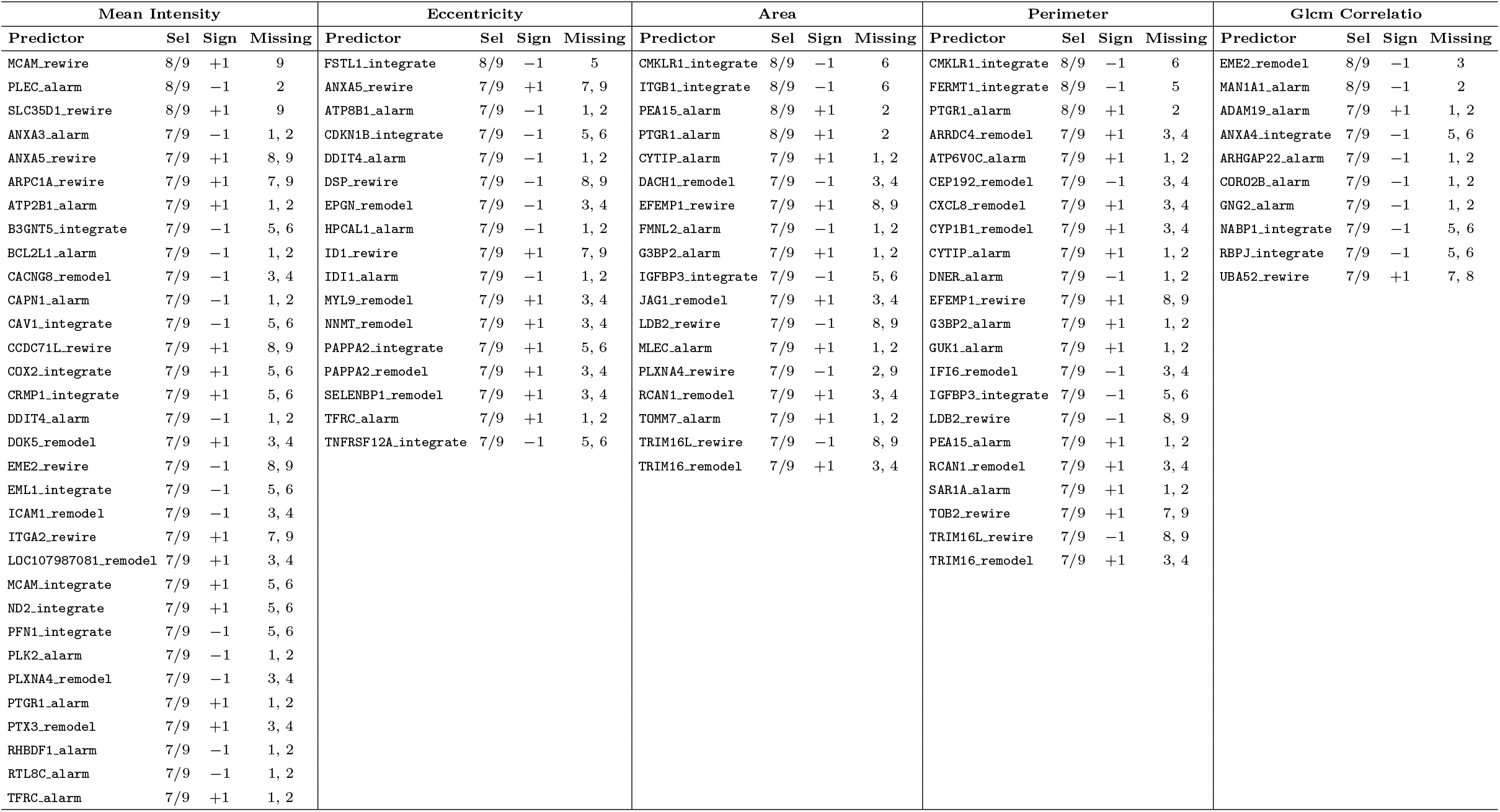
Stability-ranked phase-local gene predictors (threshold: Select_Freq ≥ 7/9). “Sel” is selection frequency across nine LOWO folds; “Sign” is the sign-consistency score; “Missing” lists held-out week(s) in which the predictor was not selected.

To better interpret folds in which an otherwise stable predictor was not selected, we recorded the held-out weeks responsible for each drop-out and summarized them alongside the stability statistics (Table 2). A consistent pattern emerged: when a phase-local gene predictor was missing in one fold, the held-out week almost always belonged to the same temporal phase as that predictor. This behavior is intuitive under the phase-local design. Because phase-local predictors are identically zero outside their phase by construction, most of the signal for estimating a phase-local coefficient comes from observations within that phase. Holding out a week from the same phase as the predictor therefore removes a nontrivial fraction of the phase-specific support—particularly in phases spanning only two weeks (alarm, remodel, integrate), where leaving out one week removes half of the within-phase data. Consequently, the evidence for selecting a phase-local predictor can weaken precisely when a week from its own phase is held out, yielding selection frequencies of 8/9 and 7/9 even for otherwise robust predictors.

Overall, these results indicate that stability ranking isolates reproducible, directionally consistent phase-local transcriptomic associations for each predictable morphology feature, while the observed missing-week pattern aligns with the intended phase-local modeling constraint rather than suggesting instability across unrelated time periods.

### 3.3 Final OLS models provide interpretable effects and enable pruning

For each morphology feature with stable transcriptomic predictors, we fit a final parsimonious OLS model including the ordinal dose covariate and the selected phase-local gene predictors, reporting HC3 robust standard errors (Section 2). Because phase-local gene predictors can be highly correlated, we applied an explicit pruning procedure: rank-deficiency resolution, VIF-based filtering to enforce VIF < 10, and removal of weakly supported predictors (e.g., *p* > 0.10) with confirmation via week-held-out performance comparisons. In several cases, removing predictors with marginal statistical support produced negligible or negative changes in LOWO *R*^2^, supporting a more compact final model. Conversely, certain predictors contributed materially to out-of-week generalization, as verified by LOWO *R*^2^ comparisons with and without the predictor.

Finally, as a parsimony check, we reassessed auxiliary time covariate (e.g., week-ordinal predictor) after including dose and phase-local gene predictors. When not supported (e.g., *p* > 0.10), week-ordinal covariate was excluded, consistent with the interpretation that remaining time-dependent structure beyond the baseline dose trend is captured by the phase-local transcriptomic effects.

#### 3.3.1 Linear model adequacy and residual diagnostics

To support interpretability-focused inference, we assess whether the final OLS models provide an adequate approximation for each morphology feature using standard regression diagnostics (Figure 5). The left column (a.1–a.5) provides plots of residuals against fitted values, while the right column (b.1–b.5) compares fitted versus observed responses relative to the identity line.

**Figure 5.**
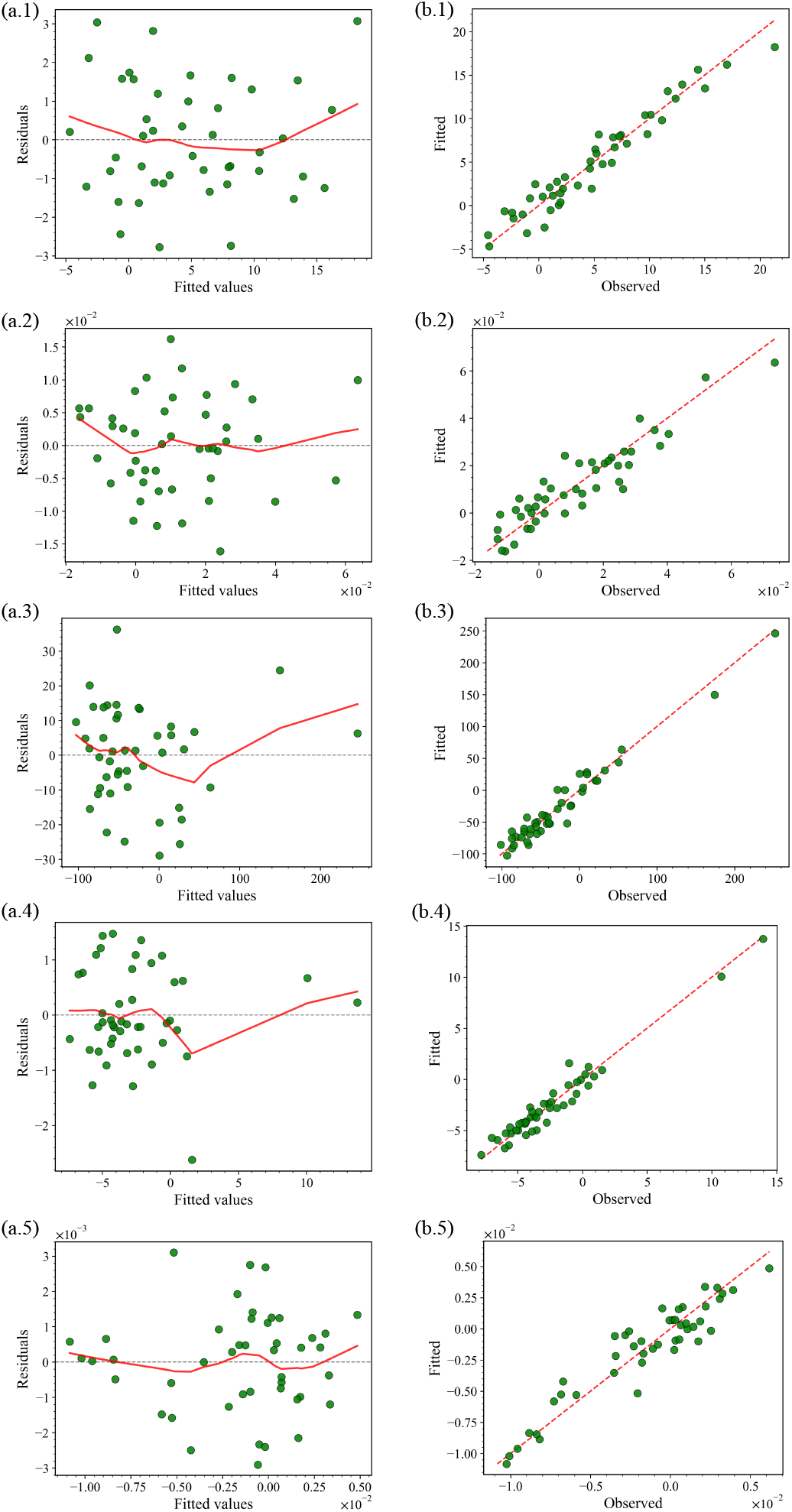
OLS model diagnostic checks across nuclear morphology features. Rows correspond (top-to-bottom) to `mean intensity, eccentricity, area, perimeter`, and `glcm correlation`. Left column (a.1–a.5): residuals versus fitted values with a smooth trend line (red) and horizontal zero reference (gray dashed). Right column (b.1–b.5): fitted versus observed values with the identity line (red dashed). Across features, residuals remain broadly centered near zero and fitted values track observations closely, supporting the adequacy of the linear model for predicting morphology features using dose and phase-local gene predictors; HC3 robust standard errors are used throughout to mitigate sensitivity to heteroskedasticity and leverage in the small-*n* setting.

##### Residual behavior

Across all five features, residuals are generally centered around zero with no pervasive, strongly directional structure, suggesting that the linear specification captures the dominant signal in each response. Several panels exhibit mild curvature in the smoothed residual trend (notably for `mean intensity`, `area`, and `perimeter`), indicating small departures from strict linearity at the extremes. We do not observe pervasive variance patterns across features that would indicate severe, systematic heteroskedasticity; however, the spread of residuals varies modestly with fitted values for some morphology features. Given the small sample size and the possibility of leverage effects from particular week–dose conditions, we report HC3 robust standard errors to stabilize inference under these modest departures from ideal OLS regression assumptions.

##### Fitted-versus-observed alignment

For each feature (Figure 5 b.1–b.5), fitted values fall close to the identity line over the observed range, indicating that the final models provide a good global approximation without obvious nonlinear distortion. The tight alignment is strongest for `mean intensity, area`, and `perimeter`, while `eccentricity` and `glcm correlation` show slightly larger relative scatter consistent with their smaller range of values. Overall, these diagnostics support the use of linear models for reporting parsimonious, phase-local gene predictors together with ordinal dose predictor, while HC3 serving as a conservative safeguard for robust standard errors.

#### 3.3.2 OLS regression summaries across morphology features

We summarize the final model behavior across the five morphology features retained for interpretable linear regression: `mean intensity, eccentricity, area, perimeter`, and `glcm correlation`. For each feature, we compare observed and fitted values as week-wise trajectories stratified by dose (Figures 6–10). Across features, the fitted curves closely track the dominant week-to-week structure within each dose condition, indicating that the final parsimonious models capture both (i) the overall exposure trend encoded by `dose_ord` and (ii) additional structured variation captured by a small set of phase-local transcriptomic predictors.

**Figure 6.**
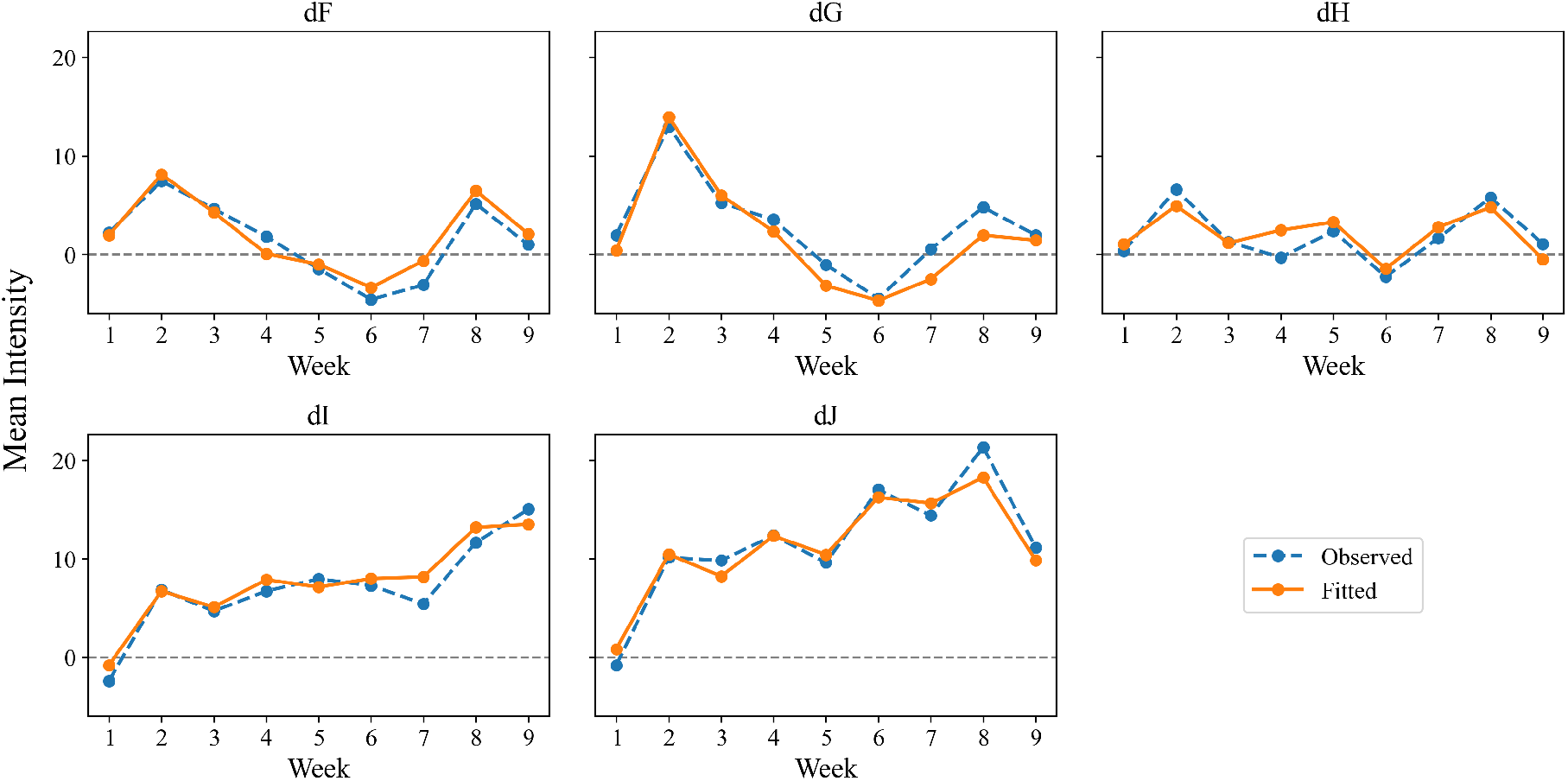
Dose-stratified observed versus fitted trajectories for mean intensity. Observed (dashed) and fitted (solid) mean intensity differences-from-control are shown across weeks for each non-control dose.

For `mean intensity`, the model reproduces the dose-stratified temporal trend across weeks with good agreement between observed and fitted trajectories (Figure 6). The corresponding OLS(HC3) fit explains a substantial fraction of variance (*R*^2^ = 0.938, adj. *R*^2^ = 0.920), and the coefficient summary (Table 3) reflects a positive `dose_ord` association alongside multiple phase-local gene effects under robust inference.

**Table 3:** OLS regression results for mean intensity (*n* = 45, *R*^2^ = 0.938, adj. *R*^2^ = 0.920). Reported standard errors are HC3-robust.

| Predictor | Coefficient | Std. Error | z-statistic | p-value | 2.5% | 97.5% |
| --- | --- | --- | --- | --- | --- | --- |
| Intercept | -2.2678 | 0.698 | -3.251 | 0.001 | -3.635 | -0.900 |
| dose_ord | 1.4902 | 0.201 | 7.402 | <0.001 | 1.096 | 1.885 |
| MCAM_rewire | 6.7755 | 2.083 | 3.254 | 0.001 | 2.694 | 10.857 |
| SLC35D1_rewire | 24.8875 | 12.544 | 1.984 | 0.047 | 0.302 | 49.473 |
| B3GNT5_integrate | -9.4165 | 2.299 | -4.095 | <0.001 | -13.923 | -4.910 |
| BCL2L1_alarm | -25.1715 | 7.019 | -3.586 | <0.001 | -38.928 | -11.415 |
| DDIT4_alarm | -6.2924 | 2.824 | -2.228 | 0.026 | -11.827 | -0.757 |
| EME2_rewire | -21.7808 | 5.854 | -3.721 | <0.001 | -33.254 | -10.308 |
| MCAM_integrate | 3.7670 | 0.813 | 4.633 | <0.001 | 2.174 | 5.360 |
| PTGR1_alarm | 14.2779 | 3.269 | 4.368 | <0.001 | 7.871 | 20.684 |
| PTX3_remodel | 9.7565 | 1.082 | 9.018 | <0.001 | 7.636 | 11.877 |

For `eccentricity`, fitted trajectories again follow the dose-specific week patterns, albeit with comparatively smaller dynamic range (Figure 7). Consistent with this, the model achieves moderate-to-strong explanatory power (*R*^2^ = 0.854, adj. *R*^2^ = 0.817) with a compact set of predictors (Table 4).

**Table 4:** OLS regression results for eccentricity (*n* = 45, *R*^2^ = 0.854, adj. *R*^2^ = 0.817). Reported standard errors are HC3-robust.

| Predictor | Coefficient | Std. Error | z-statistic | p-value | 2.5% | 97.5% |
| --- | --- | --- | --- | --- | --- | --- |
| Intercept | -0.0090 | 0.003 | -2.881 | 0.004 | -0.015 | -0.003 |
| dose_ord | 0.0043 | 0.001 | 4.251 | <0.001 | 0.002 | 0.006 |
| ANXA5_rewire | 0.1089 | 0.059 | 1.851 | 0.064 | -0.006 | 0.224 |
| EPGN_remodel | -0.0072 | 0.002 | -2.994 | 0.003 | -0.012 | -0.002 |
| HPCAL1_alarm | -0.1127 | 0.021 | -5.251 | <0.001 | -0.155 | -0.071 |
| ID1_rewire | 0.0360 | 0.008 | 4.732 | <0.001 | 0.021 | 0.051 |
| MYL9_remodel | 0.0240 | 0.007 | 3.693 | <0.001 | 0.011 | 0.037 |
| PAPPA2_remodel | 0.0154 | 0.003 | 5.544 | <0.001 | 0.010 | 0.021 |
| TFRC_alarm | 0.0701 | 0.017 | 4.063 | <0.001 | 0.036 | 0.104 |
| TNFRSF12A_integrate | -0.0591 | 0.007 | -8.204 | <0.001 | -0.073 | -0.045 |

**Figure 7.**
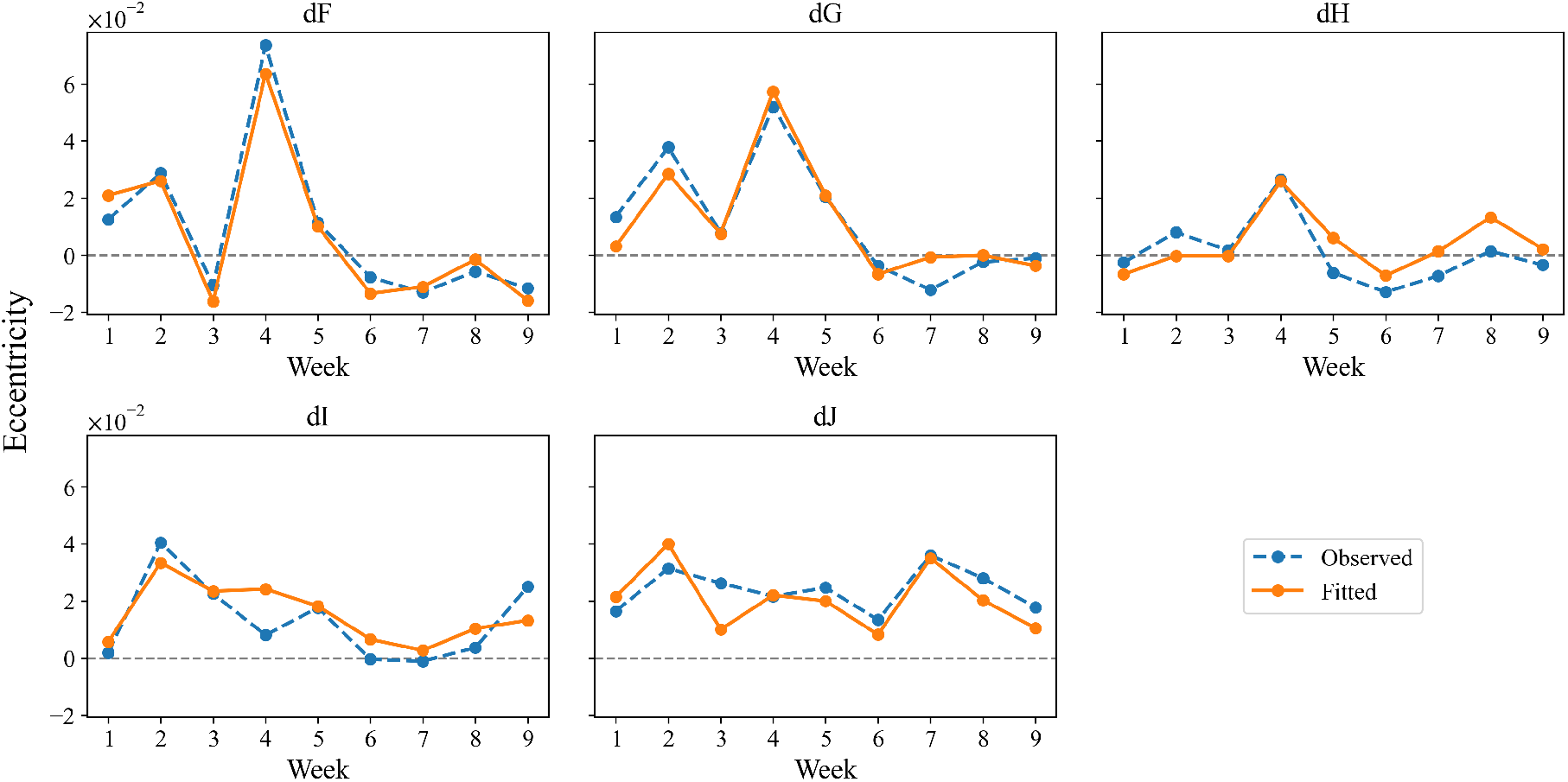
Dose-stratified observed versus fitted trajectories for eccentricity. Observed (dashed) and fitted (solid) eccentricity differences-from-control are shown across weeks for each non-control dose.

For `area` and `perimeter`, the fitted trajectories closely match observed dose-stratified trends across the full study timeline (Figures 8 and 9). These two features show the strongest overall fit among the models considered (*R*^2^ = 0.955 and 0.956, respectively; both adj. *R*^2^ = 0.944), and their OLS(HC3) summaries (Tables 5 and 6) indicate that the final retained phase-local gene predictors, together with `dose_ord`, account for the majority of observed week–dose variability. Notably, the `area` and `perimeter` models share four phase-local gene predictors (`CYTIP_alarm, PEA15 alarm, EFEMP1_rewire`, and `RCAN1_remodel`), which is expected given the strong correlation between these two morphology features and suggests coherent feature selection across related morphology features.

**Table 5:** OLS regression results for area (*n* = 45, *R*^2^ = 0.955, adj. *R*^2^ = 0.944). Reported standard errors are HC3-robust.

| Predictor | Coefficient | Std. Error | z-statistic | p-value | 2.5% | 97.5% |
| --- | --- | --- | --- | --- | --- | --- |
| Intercept | -80.7464 | 9.135 | -8.840 | <0.001 | -98.650 | -62.843 |
| dose_ord | 10.1969 | 2.653 | 3.844 | <0.001 | 4.997 | 15.396 |
| ITGB1_integrate | -125.5558 | 25.141 | -4.994 | <0.001 | -174.832 | -76.280 |
| PEA15_alarm | 218.3888 | 110.000 | 1.985 | 0.047 | 2.793 | 433.984 |
| CYTIP_alarm | 87.4482 | 11.688 | 7.482 | <0.001 | 64.540 | 110.356 |
| DACH1_remodel | -106.2618 | 26.849 | -3.958 | <0.001 | -158.884 | -53.639 |
| EFEMP1_rewire | 52.4992 | 10.179 | 5.157 | <0.001 | 32.548 | 72.450 |
| RCAN1_remodel | 219.5646 | 54.746 | 4.011 | <0.001 | 112.265 | 326.865 |
| TRIM16L_rewire | -63.3489 | 14.646 | -4.325 | <0.001 | -92.055 | -34.643 |
| TRIM16_remodel | 85.8404 | 21.306 | 4.029 | <0.001 | 44.081 | 127.600 |

**Table 6:** OLS regression results for perimeter (*n* = 45, *R*^2^ = 0.956, adj. *R*^2^ = 0.944). Reported standard errors are HC3-robust.

| Predictor | Coefficient | Std. Error | z-statistic | p-value | 2.5% | 97.5% |
| --- | --- | --- | --- | --- | --- | --- |
| Intercept | -4.2084 | 0.342 | -12.297 | <0.001 | -4.879 | -3.538 |
| dose_ord | 0.5499 | 0.175 | 3.140 | 0.002 | 0.207 | 0.893 |
| CMKLR1_integrate | -3.3494 | 0.521 | -6.432 | <0.001 | -4.370 | -2.329 |
| CEP192_remodel | -8.7338 | 1.612 | -5.418 | <0.001 | -11.893 | -5.574 |
| CYTIP_alarm | 5.9298 | 1.065 | 5.568 | <0.001 | 3.842 | 8.017 |
| DNER_alarm | -11.3347 | 3.720 | -3.047 | 0.002 | -18.625 | -4.044 |
| EFEMP1_rewire | 4.7859 | 0.892 | 5.368 | <0.001 | 3.039 | 6.533 |
| LDB2_rewire | -1.6499 | 0.521 | -3.166 | 0.002 | -2.671 | -0.629 |
| PEA15_alarm | 15.7976 | 5.530 | 2.857 | 0.004 | 4.960 | 26.635 |
| RCAN1_remodel | 9.0088 | 2.488 | 3.620 | <0.001 | 4.132 | 13.886 |

**Figure 8.**
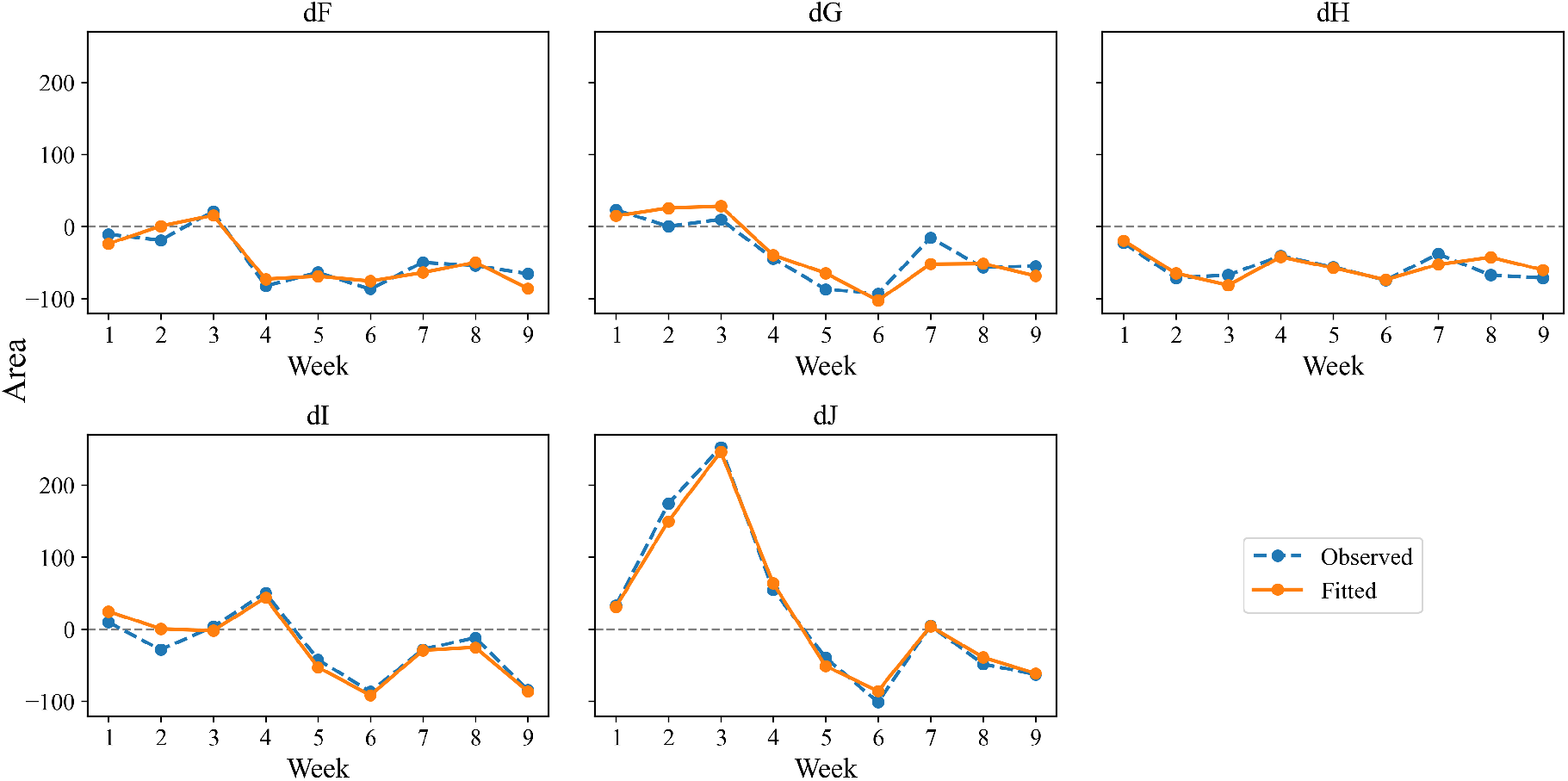
Dose-stratified observed versus fitted trajectories for area. Observed (dashed) and fitted (solid) area differences-from-control are shown across weeks for each non-control dose.

**Figure 9.**
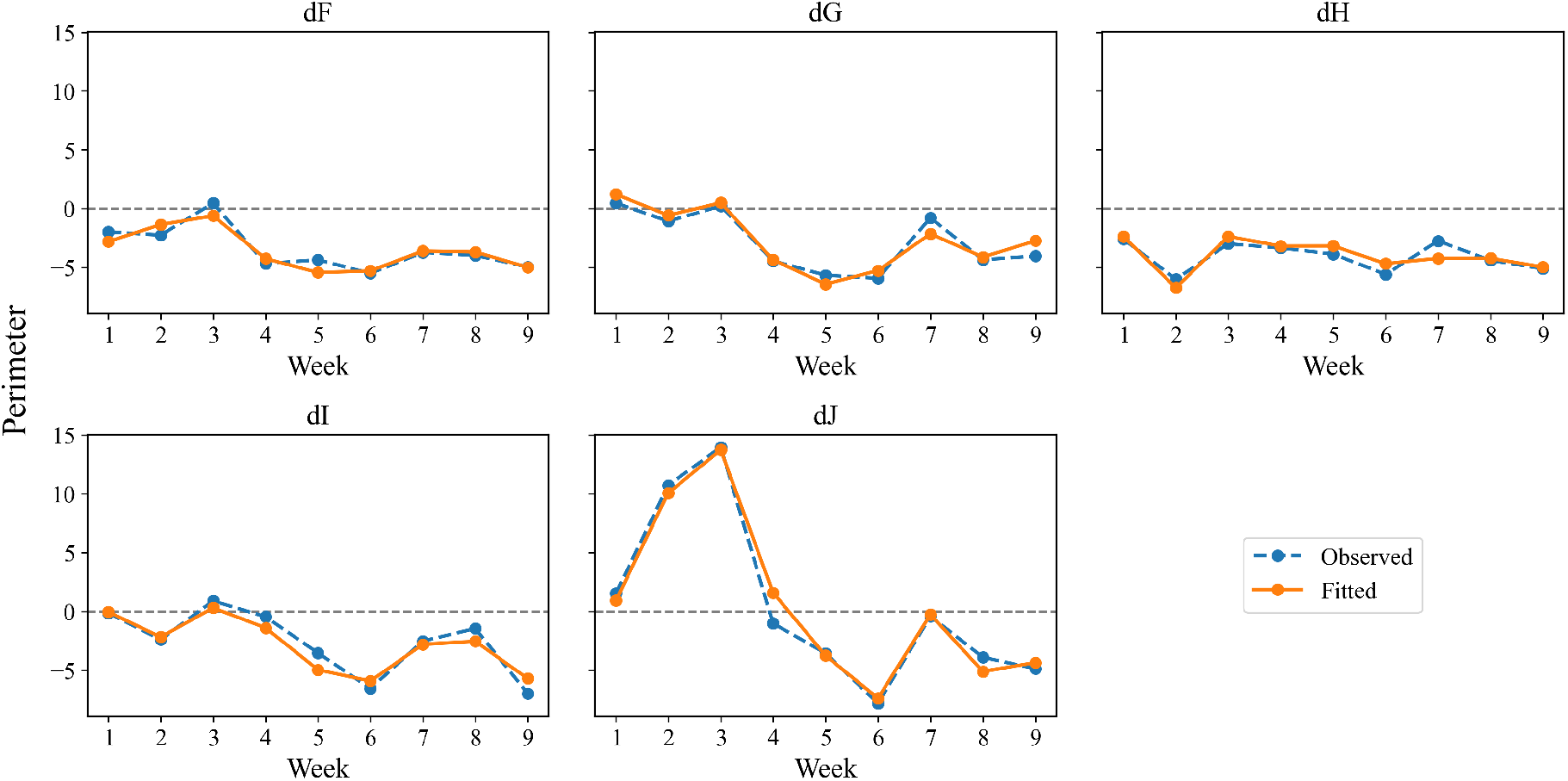
Dose-stratified observed versus fitted trajectories for perimeter. Observed (dashed) and fitted (solid) perimeter differences-from-control are shown across weeks for each non-control dose.

Finally, `glcm correlation` exhibits clear dose-stratified temporal structure that is well tracked by the fitted trajectories (Figure 10). The final OLS(HC3) model attains strong explanatory power (*R*^2^ = 0.887, adj. *R*^2^ = 0.869) with a small number of predictors (Table 7), supporting the view that a parsimonious set of phase-local transcriptomic predictors is sufficient to reproduce much of the observed week–dose variation for this texture-derived feature.

**Table 7:**
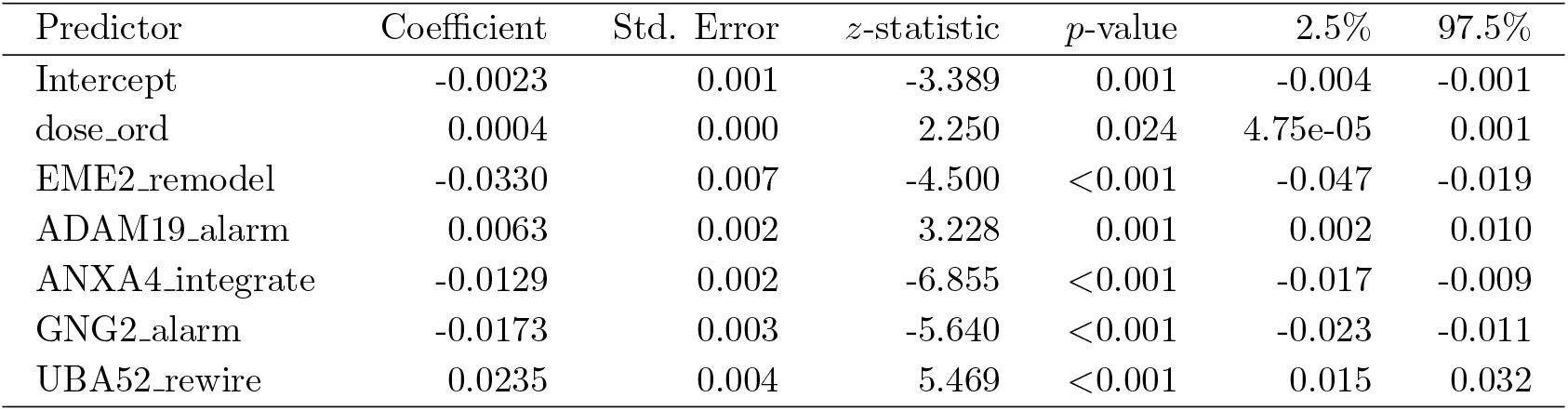
OLS regression results for glcm correlation (*n* = 45, *R*^2^ = 0.887, adj. *R*^2^ = 0.869). Reported standard errors are HC3-robust.

| Predictor | Coefficient | Std. Error | z-statistic | p-value | 2.5% | 97.5% |
| --- | --- | --- | --- | --- | --- | --- |
| Intercept | -0.0023 | 0.001 | -3.389 | 0.001 | -0.004 | -0.001 |
| dose_ord | 0.0004 | 0.000 | 2.250 | 0.024 | 4.75e-05 | 0.001 |
| EME2_remodel | -0.0330 | 0.007 | -4.500 | <0.001 | -0.047 | -0.019 |
| ADAM19_alarm | 0.0063 | 0.002 | 3.228 | 0.001 | 0.002 | 0.010 |
| ANXA4_integrate | -0.0129 | 0.002 | -6.855 | <0.001 | -0.017 | -0.009 |
| GNG2_alarm | -0.0173 | 0.003 | -5.640 | <0.001 | -0.023 | -0.011 |
| UBA52_rewire | 0.0235 | 0.004 | 5.469 | <0.001 | 0.015 | 0.032 |

**Figure 10.**
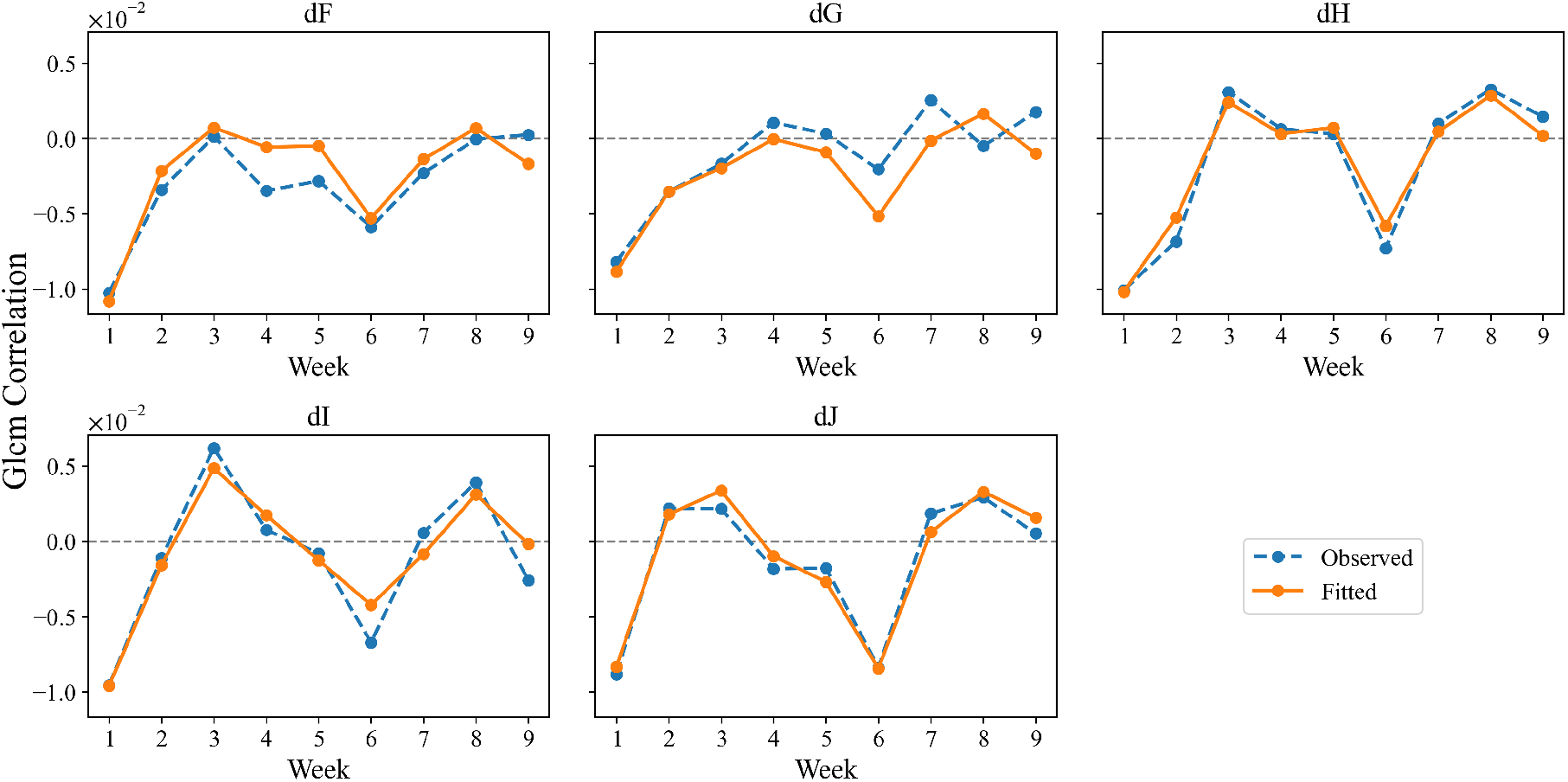
Dose-stratified observed versus fitted trajectories for glcm correlation. Observed (dashed) and fitted (solid) glcm correlation differences-from-control are shown across weeks for each non-control dose.

Across all features, we emphasize that these coefficient summaries quantify conditional, phase-local associations (given `dose_ord`) and are reported here primarily as an interpretable statistical description of the fitted models. Because predictors are selected through global screening, stability ranking, and pruning, the reported *p*-values are descriptive and conditional on the selected model rather than formal post-selection inference. Biological interpretation, relevance, and pathway-level contextualization are deferred to the Discussion section (Section 4).

## 4 Discussion

A central contribution of this work is an interpretable transcriptome-to-phenotype modeling workflow that is explicitly aligned with the longitudinal design of the experiment. Rather than relying on random train– test splits, we evaluate generalization using leave-one-week-out (LOWO) cross-validation, which stress-tests whether transcriptomic predictors can explain morphology in an unseen week. This choice is motivated by the strong week-to-week shifts observed in both modalities (e.g., culture drift, batch effects, and imaging variability), and therefore provides a conservative assessment of robustness.

Our modeling pipeline explicitly separates dominant exposure structure from additional structured variation potentially attributable to transcriptional changes. For each morphology feature, we first fit a baseline dose-only model within each LOWO fold and compute out-of-week residuals. We then fit sparse, phase-local transcriptomic models to predict these residuals. This residualization step is crucial: it reduces the risk that the transcriptomic model merely re-encodes dose through correlated gene responses, and instead prioritizes signals that improve prediction beyond a parsimonious dose predictor. Consequently, any retained phase-local predictors should be interpreted as associations with morphology conditional on the baseline dose trend.

The phase-local gene predictor design provides a compromise between a single global gene effect and a fully week-specific model. By interacting gene responses with phase indicators, the model allows the same gene to be active in some intervals and inactive in others, which is consistent with the expectation that different transcriptional programs may dominate early versus late response regimes. Importantly, this representation maintains interpretability: each coefficient corresponds to an association between a gene response and a morphology feature within a specified temporal phase. We emphasize that phase names are shorthand descriptors of contiguous week ranges and do not, by themselves, assert pathway identity.

We use stability ranking to identify phase-local predictors that recur across LOWO fits, summarizing selection frequency and sign consistency. In this longitudinal setting, stability values such as 7/9 or 8/9 can still be informative: phases spanning only two weeks provide limited within-phase support, and holding out one week can remove a substantial fraction of the phase-specific evidence. Observing that some predictors are absent primarily when a week from their own phase is held out is therefore not necessarily pathological; instead it may indicate that the association is genuinely phase-local and depends on information contained in that time window. Nevertheless, these patterns can also flag weeks that behave differently (e.g., technical anomalies) and merit closer scrutiny.

The final OLS stage is primarily a reporting layer: after selecting a compact set of stable predictors and pruning multicollinearity, we fit an interpretable linear model and report coefficients with heteroskedasticity-consistent (HC3) standard errors. This provides conventional confidence intervals and readable effect-size summaries while acknowledging that upstream selection occurred. The use of robust (HC3) covariance is intended to reduce sensitivity to leverage points and modest heteroskedasticity that can arise in small-*n* regression, particularly when predictors are partially collinear or when the response variance varies across dose levels.

To illustrate the biological interpretability afforded by the phase-local framework, we examine the predictors selected for nuclear mean intensity, which achieved the strongest residual predictability and highest final model fit (*R*^2^ = 0.938). The alarm-phase predictors *BCL2L1* and *DDIT4* encode two canonical early stress-response proteins: *BCL2L1*, whose major isoform BCL-xL is an anti-apoptotic member of the BCL-2 family that inhibits radiation-induced apoptosis by preventing mitochondrial cytochrome c release (Kharbanda et al., 1997), and *DDIT4* (also known as REDD1), a p53-dependent mTORC1 inhibitor that limits the apoptotic response following ionizing radiation through feedback control of p53 translation (Vadysirisack et al., 2011). The remodel-phase predictor *PTX3* (*p*-value ≈ 1.92 × 10^−19^), the single most significant gene predictor across all morphology features, encodes a long pentraxin that functions as a pattern recognition molecule in innate immunity and plays direct roles in extracellular matrix remodeling through interactions with fibrinogen, collagen, and hyaluronic acid (Garlanda et al., 2018). Critically, PTX3 has been shown to be persistently upregulated in irradiated human arteries and veins compared to matched non-irradiated controls (Christersdottir Björklund et al., 2013), establishing it as a marker of sustained radiation-induced tissue damage. In the rewire phase, *EME2*, encoding the non-catalytic subunit of the MUS81-EME2 structure-selective endonuclease that promotes replication fork restart through nucleolytic cleavage of stalled forks (Pepe and West, 2014), shows a negative association consistent with resolution of persistent replication stress in the later weeks. Together, these predictors offer a biologically plausible interpretation of the temporal trajectory of nuclear mean intensity (Figure 2): early pro-survival and growth-suppressive signaling (alarm) correlates with the initial elevation of nuclear intensity across all dose groups, sustained innate immune activation and matrix remodeling (remodel) coincides with the onset of dose-dependent divergence at intermediate weeks, and declining replication fork rescue activity (rewire) aligns with the sustained separation of higher dose groups in the later weeks. No integrate-phase predictor was retained for this feature, suggesting a gradual transition between the remodel and rewire programs rather than a molecularly distinct intermediate step. At higher dose rates, particularly doses I and J, the stronger and more sustained elevation of intensity is consistent with prolonged activation of these transcriptional programs.

The area and perimeter models (both *R*^2^ ≈ 0.955) share four phase-local predictors whose concordant selection across these highly correlated yet independently modeled features serves as an internal consistency check. The normalized morphological trajectories reveal a striking transient enlargement at higher dose rates—most pronounced at dose J during weeks 2–3—followed by a contraction to or below control levels by mid-experiment. In the alarm phase, *PEA15*, an anti-apoptotic protein that sequesters ERK/MAPK in the cytoplasm (Formstecher et al., 2001), shows a positive association with nuclear size, correlating with the early transient enlargement observed at higher dose rates. The remodel-phase predictor *RCAN1*, an endogenous inhibitor of calcineurin/NFAT signalling (Lao et al., 2022), coincides with the peak and subsequent decline of nuclear area. The area model additionally retains *ITGB1* in the integrate phase—encoding integrin beta-1, a key mediator of cell–extracellular matrix adhesion (Su et al., 2024) bridging the remodel and rewire programs, though this predictor is not shared with perimeter. In the rewire phase, *EFEMP1* (fibulin-3), a secreted ECM glycoprotein that modulates matrix integrity through regulation of metalloproteinase activity (Livingstone et al., 2020), aligns with the later weeks when nuclear dimensions have contracted below control levels, implicating extracellular matrix restructuring in the long-term morphological outcome. The dose-dependent magnitude of the transient enlargement, particularly at dose J, is consistent with stronger activation of these programs at higher dose rates. While these associations are predictive rather than causal, the recovery of genes with established roles in radiation response, oxidative stress signaling, and tissue repair—spanning temporally distinct phases and recurring across correlated morphology features—provides evidence that the phase-local modeling framework identifies biologically interpretable transcriptome–phenotype relationships suitable for downstream validation.

Several limitations should be stated explicitly. First, the reported coefficients represent predictive associations, not causal mechanisms; the gene-level interpretations offered above for nuclear mean intensity and area/perimeter are illustrative examples drawn from individual predictors rather than a comprehensive biological account. Second, the candidate-gene screening procedure and phase definitions constrain discovery to response-enriched subsets and may omit relevant genes that are weakly responsive by differential expression but still important for morphology. Third, the modeling assumes a linear additive structure within each phase; non-linearities and interactions may exist but are not captured in the current interpretable specification. Finally, statistical support in this framework should be viewed as prioritization for follow-up rather than definitive biological validation.

A natural next step is to move from individual gene interpretations presented here to pathway-level interpretation, for example by performing phase-wise enrichment analyses on stable predictor sets and comparing enriched pathways across phases and across morphology features. Such an approach would generalize the gene-specific observations (e.g., the alarm-phase stress and anti-apoptotic signatures, the remodel-phase innate immune and matrix remodeling programs) into broader functional themes. Orthogonal validation could include aligning inferred phases with independent readouts (e.g., cell-cycle profiling, DNA damage markers, chromatin accessibility, or senescence-associated markers) and testing whether perturbations of candidate genes shift nuclear morphology in predicted directions. These extensions would strengthen the biological grounding of the transcriptome-to-phenotype links while preserving the interpretability and week-held-out robustness emphasized here.

## 5 Conclusion

In this study we have developed a transparent, week-held-out modeling framework to relate RNA-seq responses to Cell Painting-derived nuclear morphology features under chronic low-dose radiation exposure. By expressing both modalities as within-week differences relative to matched controls and evaluating generalization with leave-one-week-out cross-validation, the pipeline directly targets transcriptome–phenotype relationships that persist beyond week-to-week shifts. The core methodological choices—dose-only residualization, phase-aware gene–time interactions, sparse Elastic Net fitting with an explicit sparsity cap, and stability-based ranking—provide a practical route to interpretable candidate predictors in a small-*n*, high-dimensional longitudinal setting.

Across multiple nuclear morphology features, the final parsimonious linear regression models achieved strong goodness-of-fit while maintaining interpretability through phase-dependent coefficient summaries. Dose-stratified trajectory comparisons indicate that the selected predictors can reproduce the dominant temporal patterns within dose conditions rather than only fitting aggregate trends. Importantly, the identified phase-aware gene predictors should be interpreted as predictive, conditional associations that prioritize hypotheses for follow-up, not as mechanistic or causal claims. Future work will extend interpretation to pathway-level summaries, assess sensitivity to alternative phase definitions and screening thresholds, and validate top predictors using targeted perturbations and independent readouts.

## Acknowledgment

This research was funded by the Biological and Environmental Research program in the US DOE’s Office of Science under project B&R# KP1601017 and FWP#CC140.

## A Elastic Net hyperparameter grid search and solver settings

For each nuclear morphology feature, Elastic Net regression hyperparameters are tuned using leave-one-week-out (LOWO) cross-validation. Within each fold, phase-local predictors are standardized using statistics computed on the training weeks only, and the same affine transform is applied to the held-out week. To handle the intercept consistently across solvers, training residuals are mean-centered and the training-fold mean is added back to predictions on the held-out week.

We sweep a grid over the global regularization strength *α* and the mixing parameter *ρ* (“*ℓ*_1_-ratio”) which mixes (*ℓ*_1_, *ℓ*_2_) penalties. The primary grid uses a log-spaced range of *α* values spanning weak to strong regularization as follows:

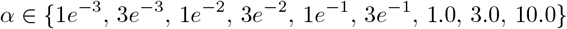

wherein this range is expanded toward larger *α* if needed to obtain sufficiently sparse model fits, and a fixed set of *ρ* values,

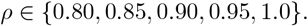

For computational efficiency, for each fold and each *ρ* we compute the full solution path across all *α* values using the coordinate-descent path routine (`sklearn.linear_model.enet_path`). The *α* values are supplied in decreasing order to exploit warm starts in the path solver. Unless stated otherwise, path computations use a conservative iteration budget and tolerance (default: `max_iter=200000, tol=`10^−5^).

Out-of-fold residual predictions are aggregated across folds, and each (*α, ρ*) setting is scored by the correlation 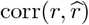. Hyperparameters are selected by maximizing this score subject to a per-fold sparsity cap on the number of nonzero coefficients (default cap: max(nnz) ≤ 50); if no setting satisfies the cap, the overall best-scoring setting is used.

For stability ranking at the selected (*α, ρ*), we refit Elastic Net regression model within each training fold using `sklearn.linear_model.ElasticNet` with `fit_intercept=False`, deterministic coordinate updates (`selection=“cyclic”`), `max_iter=50000,` and tolerance `tol=`10^−5^.

## Notes

### Competing Interest Statement

The authors have declared no competing interest.

